# Replication timing alterations in leukemia reflect stable clinically-relevant changes in genome architecture

**DOI:** 10.1101/549196

**Authors:** Juan Carlos Rivera-Mulia, Takayo Sasaki, Claudia Trevilla-Garcia, Naoto Nakamichi, David Knapp, Colin Hammond, Bill Chang, Jeffrey W. Tyner, Meenakshi Devidas, Jared Zimmerman, Kyle N. Klein, Vivek Somasundaram, Brian Druker, Tanja A. Gruber, Amnon Koren, Connie Eaves, David M. Gilbert

## Abstract

Human B-lineage precursor acute lymphoid leukemias (BCP-ALLs) comprise a group of genetically and clinically distinct disease entities with features of differentiation arrest at known stages of normal B-lineage differentiation. We previously showed BCP-ALL cells display unique and clonally heritable DNA-replication timing (RT) programs; i.e., programs describing the variable order of replication and sub-nuclear 3D architecture of megabase-scale chromosomal units of DNA in different cell types. To determine the extent to which BCP-ALL RT programs mirror or deviate from specific stages of normal human B-cell differentiation, we transplanted immunodeficient mice with quiescent normal human CD34+ cord blood cells and obtained RT signatures of the regenerating B-lineage populations. We then compared these with RT signatures for leukemic cells from a large cohort of BCP-ALL patients of varied genetic subtype and outcome. The results identify BCP-ALL subtype-specific features that resemble specific stages of B-cell differentiation and features that appear associated with relapse. These results suggest the genesis of BCP-ALL involves alterations in RT that reflect biologically significant and clinically relevant leukemia-specific epigenetic changes that have potential as a novel genre of prognostic biomarkers.

**KEY POINTS:**

- DNA replication timing of >100 pediatric leukemic samples identified BCP-ALL subtype-specific genome alteration signatures.
- Comparative analysis identified features that resemble specific stages of B-cell differentiation and features associated with outcome.

## INTRODUCTION

DNA replication timing (RT) refers to the temporal order in which defined units of chromosomes replicate during the course of S phase. The regulatory units of RT correspond to units of structural organization and are organized into higher-order 3D spatial compartments in the nucleus that replicate at distinct times during S phase ^1,2^. Changes in RT affect at least half of the genome during normal development and differentiation ^1,3,4^ and RT profiles are characteristic of a given cell type ^5-8^. Early RT correlates with transcriptional activity, but there are many exceptions ^9,10^ and “RT signatures” can identify differences between diseased and normal tissue that are not identified by standard transcriptome analyses ^11,12^. RT signatures may therefore provide a novel genre of clinical biomarkers reflective of large-scale genome architecture. We previously described disease- and patient-specific features in the RT profiles of B-lineage precursor acute lymphoid leukemia (BCP-ALL) cells ^2,13^ and showed these remained stable in serially passed patient-derived xenografts (PDX) in immunodeficient mice ^14^. Here, we investigated the biological relevance of RT alterations to BCP-ALL by examining the relationship of BCP-ALL RT profiles to specific stages of normal B-cell differentiation from which this class of leukemias derive, and their potential prognostic significance. Results establish the existence of leukemia-specific RT signatures that include previously unknown associations with specific BCP-ALL subtypes and their responses to therapy.

## METHODS

### Patient Samples

Primary BCP-ALL patient samples were obtained with informed consent according to protocols approved by the Institutional Review Boards of Oregon Health & Science University and St. Jude’s Childrens Research Hospital. Mononuclear cells were obtained from bone marrow aspirates by Ficoll density gradient centrifugation, and viably frozen cells were stored in 90% fetal bovine serum, 10% DMSO.

### Normal Cells

Normal human cord blood samples were obtained with informed consent, anonymized and used according to procedures approved by the Research Ethics Board of the University of British Columbia. Low density CD3-CD19-CD11b-cells simultaneously depleted of neutrophils and red cells were isolated on Lymphoprep using RosetteSep™ and the >90% pure CD34+ cells isolated using EasySep (both STEMCELL Technologies). Cells were stored frozen at −176 °C in DMSO with 90% fetal bovine serum (FBS). Prior to use for transplanting mice the cells were thawed in Iscove’s modified Dulbecco medium with 10% FBS (both STEMCELL Technologies) and 10 mg/mL DNase I (Sigma Aldrich), centrifuged, and re-suspended in Hanks balanced salt solution (STEMCELL Technologies) with 2% FBS.

### Xenografts

2-10 × 10^4^ normal human CD34+ CB cells (2 biological replicates consisting of pooled CB cells from 3 individuals) were intravenously injected into 8-12 week adult female NRG mice within a few hours of being exposed to 8.5 cGy of ^137^Cs γ-rays delivered over 3 hours. These mice were bred in the Animal Resource Centre of the British Columbia Cancer Research Centre and treated using procedures approved by the Animal Care Committee of the University of British Columbia. 10-15 weeks later, pelvic, femoral and tibial bone marrow and spleen cells were isolated, and sorted for subsets by FACS^15^.

### Simultaneous Sorting for Surface Markers and DAPI

Red blood cell lysis was first performed using Ammonium Chloride solution from STEMCELL Technologies for 10 minutes on ice. A crude enrichment for human cells was performed using an EasySep™ Mouse/Human Chimera Isolation Kit (STEMCELL Technologies) as per manufacturer instructions but with only a single separation on the magnet and with an extra wash of the mouse cells retained on the magnet during the pour off. Cells were then spun down and resuspended in Iscove’s Modified Dulbecco’s Medium (IMDM) supplemented with 10% Fetal Bovine Serum (both from STEMCELL Technologies) and 100 µM Bromodeoxyuridine (BrdU, BD Pharmingen) and were placed in a humidified 37 °C (5% CO2 in air) for 2 hours. Following incubation, cells were washed once with Dulbecco’s phosphate-buffered saline (PBS) + 2% FBS, and stained with 1:25 anti-human CD45 FITC (clone 2D1, STEMCELL Technologies), 1:50 anti-human CD45 APC-ef780 (clone HI30, eBiosciences), 1:100 anti-human CD34 AlexaFluor 647 (clone 581), 1:50 anti-human CD19 PE-Cy7 (clone HIB19), and 1:800 anti-human ROR1 PE (clone 2A2, all from Biolegend) for 30 minutes on ice. Cells were once again washed, then resuspended in 8 mg/mL DAPI (Sigma) + 200 mg/mL Digitonin (Sigma) in PBS + 2% FBS. Cells were then sorted on a BD FACSAria™ Fusion sorter directly into Qiagen Buffer AL for downstream processing.

### Repli-chip, Repli-seq and RT signatures

Repli-ChIP ^16^ and Repli-seq ^17^ were performed as detailed elsewhere. Genome-wide RT profiles were constructed, scaled and pooled for analysis as previously described ^16,17^. RT signatures were identified by unsupervised clustering as described ^7 18 19^.

### Capture Repli-seq

Roche SeqCap EZ Developer Library (cat # 06471684001, IRN/Design Name:<u>4000016400</u>) was designed to tile a 250 bp capture region within the central 4-kb target region from each 10-kb window of hg19 avoiding non-specific sequences by Roche so that the maximum distance between 2 capture regions is 14-kb, with a 6-kb minimum distance between 2 capture regions. For Capture Repli-seq, G1 and S phase total genomic DNA libraries (average 150 bp insert size) were made from cells isolated by FACS from each patient sample using NEBNext Ultra DNA Library Prep Kit for Illumina (NEB E7370) and single indexed using NEB E7335 or NEB E7500. Up to 12 each of these indexed libraries were pooled and representative 250 bp regions from each 10-kb window throughout the genome captured using the Roche SeqCap EZ Developer Library (cat # 06471684001, IRN/Design Name:<u>4000016400</u>), the SeqCap EZ Pure Capture Bead Kit (cat #06 977 952 001), and the SEQCAP EZ REAGENT KIT PLUS (cat #06 953 212 001), according to manufacturer’s instructions. After capture, enrichment of the target region was confirmed by qPCR. The target region was 51-221 times enriched compared with the pre-capture library and non-target regions were reduced by 0.02-0.16 times of pre-capture library.

### RT Profiles from Deep WGS

Paired-end WGS on diagnostic leukemic blasts and matched germ line DNA samples from TCF3-PBX1 patients treated at St. Jude Children’s Research Hospital was performed using the Illumina sequencing platform (Illumina Inc., San Diego, CA). The leukemic genomes had an average haploid coverage of 28x. Replication timing was inferred from DNA copy number as previously described ^20^.

### TCF3-PBX1 siRNA Transfection

Cell lines were transfected using the Lonza Amaxa Nucleofector kit CA 137 program with 6 μg each of 2 separate E2A-PBX1-specific siRNAs (Eurofins/Operon): CUC CUA CAG UGU UUU GAG U and CAG UGU UUU GAG UAU CCG. These siRNA’s were tagged at the 3’end with Cyanine-3 NHS Ester (Cy-3) and Cyanine-5 NHS Ester (Cy-5), respectively, to monitor transfection efficiency by fluorescence microscopy.

## RESULTS

### RT signatures of normal human hematopoietic cell types

We transplanted normal human CD34+ CB cells into sub-lethally irradiated Non-obese diabetic-*Rag1*^-/-^ *IL2Rγc*^-/-^ (NRG) mice and isolated human CD34+CD38-, CD34+CD19+, CD34-CD19+ and total CD34+ (mixed progenitors) cells by fluorescent activated cell sorting (FACS) from their bone marrow and spleens 2-3 months later (**Figure 1A**). Biological replicates consisted of cells regenerated from different pooled sources of CB cells. The CD34+CD19+ and CD34-CD19+ cells obtained in one of the replicate experiments were also sorted for the presence or absence of *ROR1* expression. As expected ^21^, CD34+CD19+ROR1+ cells were rare allowing purification of only 300 cells from the early and late S-phase fractions, simultaneously sorted for surface marker expression and DNA content, from which good quality Repli-seq profiles were then also obtained.

**Figure 1.**
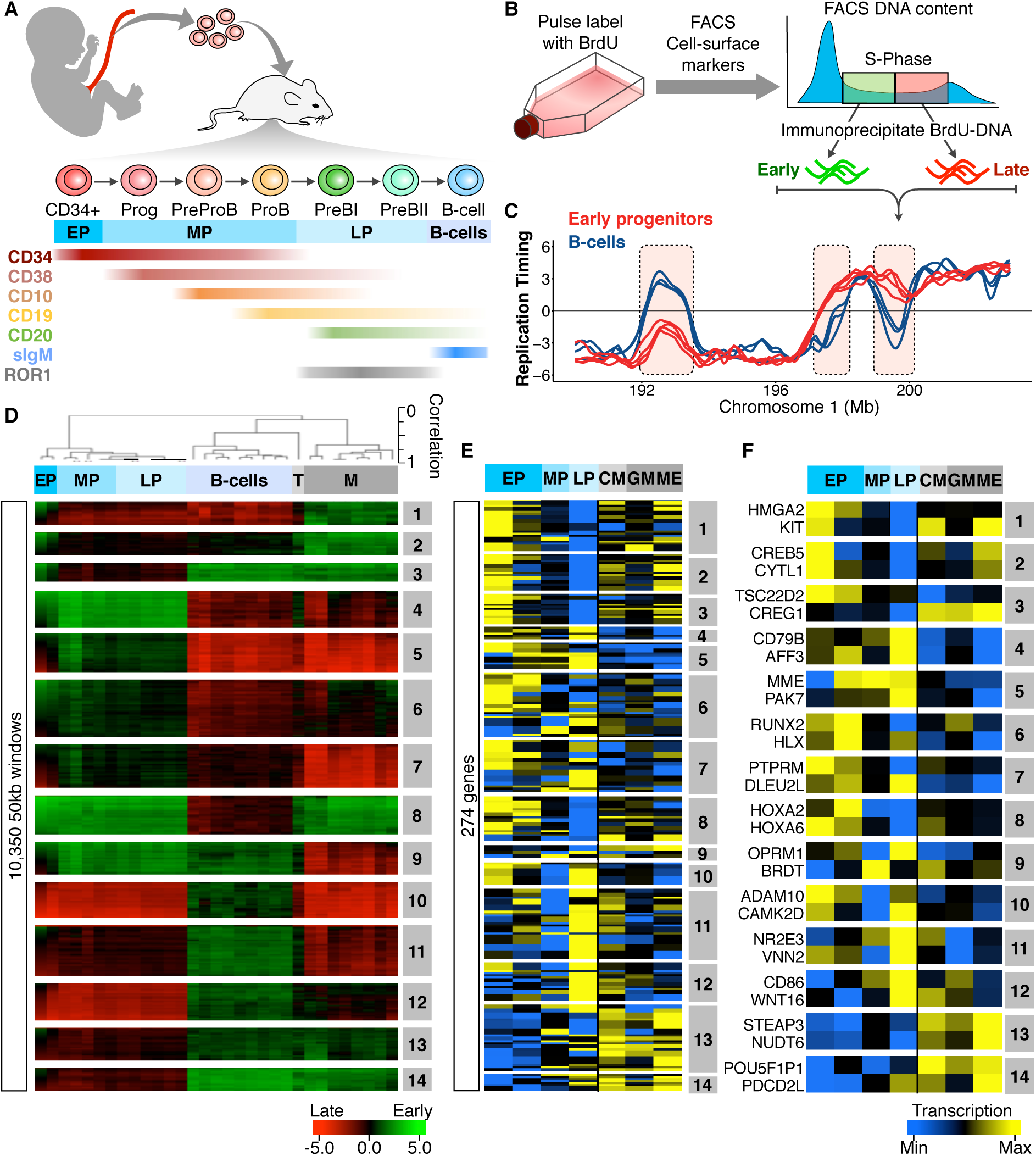
RT changes during B-cell development. **(A)** HSCs extracted from cord blood (CB) were expanded in xenografted mice and developmental intermediates of B-cell differentiation were extracted from xenografted mice by FACS using the cell surface markers shown in distinct colors. The approximate temporal expression patterns during differentiation are shown. (**B**) RT profiling method. **(C)** Exemplary RT profiles highlighting in pink differences between immature (red) and mature B-cells (blue). **(D)** Unsupervised clustering analysis of RT-variable regions identified specific RT signatures (labeled in gray boxes). Dendrogram at the top shows the correlation of all samples based on the 10,350 RT-variable 50kb segments identified. The heat map shows the RT ratios [= log2(early/late)]. (**E**) Early human hematopoiesis transcriptome data (Laurenti et al., 2013) was used to measure gene expression patterns within each RT signature. RT signatures from (D) contained 274 genes with transcriptional patterns coordinated with their RT program. (**F**) Heatmap of expression patterns for known hematopoietic regulators within the RT signatures from (D). Two exemplary genes for each RT signature are shown.

RT profiles were generated by Repli-chip ^16^ or Repli-seq ^17^ as described in the Methods and illustrated in **Figure 1B** and **1C**. RT signatures specific to differentiating B-cells were derived by comparisons with similarly generated RT profiles obtained for other normal human hematopoietic cells, including our previously reported data for granulopoietic and erythroid cells generated from adult mobilized peripheral blood cells, stimulated adult T-cells, and a series of Epstein Barr virus (EBV)-immortalized B-cell lines ^7^ with a stringent quality control cut-off ^16,17^.

We then derived RT “signatures” using a pipeline that identifies regions of the genome that replicate at times unique to each of 26 normal human cell types ^7^. In brief, after removing reads from the sex chromosomes, remaining mappable reads for each cell type were first divided into 55,940 50-kb segments each of which was then assigned an RT value (**Supplemental Figure S1A**). We then applied an unsupervised K-means clustering analysis to all 50-kb genomic segments that changed their RT from a log_2_ ratio ≥0.5 to ≤–0.5 (or vice versa) ^7^ to identify 10,350 features (18.5% of the genome) that replicate significantly differently between cell types (p-value <2×10^-16^ using T-tests). Correlation matrix (**Supplemental Figure S1B**) and dendrogram (**Figure 1D**) analyses of all samples using only these RT-variable regions confirmed the close matching of data from separately analyzed biological replicates (cells generated from different donor pools) and the clustering of samples by hematopoietic cell type to yield 14 RT signatures (**Figure 1D and Supplemental Figure S1C-D**). Interestingly, some of the RT-variable regions displayed multiple RT switches between sequential early stages of normal human B-cell differentiation. Gene expression changes for each of these signatures, derived from published human hematopoiesis transcriptome data ^22^ were generally coordinated with their RT changes (**Figure 1D-F)**. Moreover, multiple known key regulators of hematopoiesis were found in each signature; for example, *HMGA2, KIT* and *HOX* genes that are required for self-renewal activity in hematopoietic stem cells ^23,24^; *CREB5* linked to quiescence of early precursors; *RUNX2* required for HSC differentiation ^25^; and *CD79B* is required for the pro-B/pre-B transition ^26^ (**Figure 1D-F and Supplemental Table S2**). Together, these results indicate that RT features can discriminate phenotypically different hematopoietic cell types.

### BCP-ALL-specific RT signatures

Genome-wide RT profiles generated on a panel of **97** BCP-ALL patient samples and cell lines, and more limited numbers of pediatric AML and T-ALL patient samples and cell lines (**Supplemental Table S1)**. Although our standard E/L method (**Figure 2A**) performs well on samples of freshly obtained leukemic cells from patients ^13^ or transplanted mice ^14^, the numbers of viable cells or cells able to incorporate BrdU that we obtained from many frozen banked samples were very low, consistent with S-phase specific replication fork collapse ^27^. For these we first sorted unlabeled S-phase and G1 phase cells by FACS based on their DNA content (**Figure 2A**). We then used microarray or whole genome sequencing (WGS) to obtain the relative copy numbers of each 50-kb sequence in S and G1 and derived RT profiles from the fact that earlier replicating sequences are higher in copy number than late replicating sequences in S phase, but are at equal copy number in G1 phase (**Figure 2A; S/G1 Repli-seq**) ^14,28^. Since the dynamic range of G1/S data is inherently <2-fold, a large number of sequencing reads is needed to quantify copy numbers (160 million per sample), making it expensive as a routine assay. Thus, we designed a set of capture oligonucleotides spaced evenly throughout the genome (see Materials and Methods) that significantly reduced the sequence depth required to cover the breadth of the genome (10^7^ mappable reads) and also enabled high resolution RT profiles to be obtained, scaled and normalized with all other samples (**Figure 2B**). Clustering demonstrated a high concordance between replicate datasets and confirmed a lack of bias from the different methods used (**Figure 2C**).

**Figure 2.**
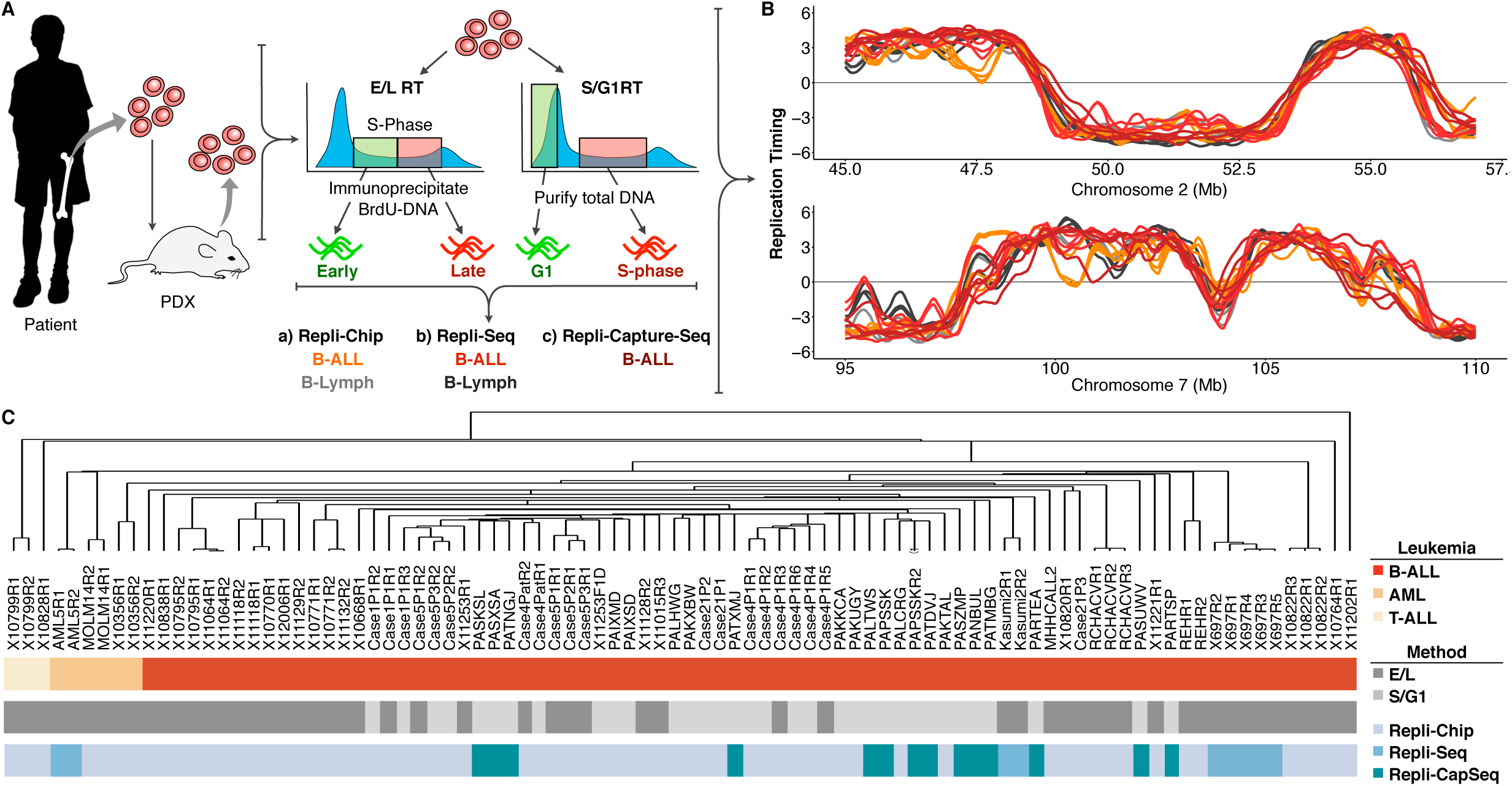
Protocols for genome-wide RT profiling. **(A)** Bone marrow samples were collected from pediatric BCP-ALL patients and analyzed by one of the methods shown. Samples with high viability were analyzed by E/L methods while samples with poor viability were either analyzed by S/G1 methods or they were expanded in patient derived xenografts (PDX) to rejuvenate their viability for analysis by E/L methods (Sasaki et al., 2017). RT was measured using either E/L or S/G1 methods (Gilbert, 2010; Sasaki et al., 2017). In the E/L method, asynchronously growing cells are pulse-labeled with 5’-bromo-2’-deoxyuridine (BrdU) and separated by FACS into early and late S phase populations. BrdU-substituted DNA is immunoprecipitated with an anti-BrdU antibody and subjected to either microarray analysis (E/L Repli-chip; Ryba et al., 2011) or NextGen sequencing (E/L Repli-seq; (Marchal et al., 2018) to obtain a replication timing ratio [=log_2_(Early/Late)] along each chromosome. For the S/G1 method, unlabeled cells are sorted into S phase and G1 phase fractions, the relative abundance of genomic sequences is quantified by either microarray (S/G1 Repli-chip) or evenly spaced segments of genomic DNA are captured using oligonucleotides and subject to NGS (Repli-CaptureSeq). **(B)** Exemplary RT profiles showing the similarity in results using the various methods after appropriate local polynomial-smoothing (Loess). **(B)** Hierarchical clustering analysis of all patient datasets based on their genome-wide RT programs confirms that there is. no bias from the different methods exploited for RT profiling.

Comparison of the RT programs from the leukemia patient samples and cell lines with one another using the same pipeline revealed 3,540 50-kb variable segments (6.3% of the autosomal genome) that showed significant differences and generated 14 RT signatures (**Figure 3A**). Six of these were linked to patients and cell lines containing the t(1;19) translocation that encodes the TCF3-PBX1 fusion protein (#1 to #6 in **Figure 3A** and **3B).** *TCF3* (also known as E2A) is a transcription factor required for normal B- (and T-) cell differentiation and has been implicated in many lymphoid malignancies ^29,30^. *PBX1* is a proto-oncogene with a critical role in hematopoiesis and lymphopoiesis ^31^. The *TCF3-PBX1* fusion gene is believed to be a driver mutation in the BCP-ALLs in which it is found ^32^. The #1 and #2 RT signatures were early replicating and #6 was late replicating uniquely in the *TCF3-PBX1*-positive samples. The fact that all 3 *TCF3-PBX1*-positive cell lines shared the same RT signatures as the patients’ *TCF3-PBX1*-positive leukemic cells underscores the epigenetic stability of their identified RT signatures. Interestingly, signatures #3, #4 and #5 sub-stratified the *TCF3-PBX1* samples, with signatures #3 and #5 found to be shared with an established cell line from an AML patient (MOLM1), but not with any of the two other AML patients’ samples. In addition, most of the regions in RT signature #1 that were early replicating uniquely in the *TCF3-PBX1*-positive patients’ cells were also very late in both T-ALL and AML cells. Commonalities between T-ALL and AML cells may not be driven by known similarities between AML and Early T-cell precursor T-ALL (ETP ALL) ^33^, since T-ALL sample 10-799 is not categorized as ETP ALL (**Supplemental Table S1**) and RT signatures #3 and #5, which were common to the AML and some *TCF3-PBX1*-positive samples, also distinguished AML from T-ALL. RT signature #8 was early replicating in a group of BCP-ALL samples of unknown genotype, and #9 was shared between these and certain AML samples. RT signatures #10, #11, and #12 identified known subsets of BCP-ALL samples. Signature #13 was late in one case and #14 was late in 2 others. The exclusivity of these RT signatures for specific subsets of patients can be further probed by clustering all patient samples using only the 50-kb chromosomal segments (features) found in each of the RT signatures. For example, clustering analysis of patient samples using the chromosomal segments from RT signature #1 identified the patients carrying the *TCF3-PBX1* translocation (**Figure 3C**). Heat maps and dendrograms for each RT signature from **Figure 3A** are shown in **Supplemental Figure S2**.

**Figure 3.**
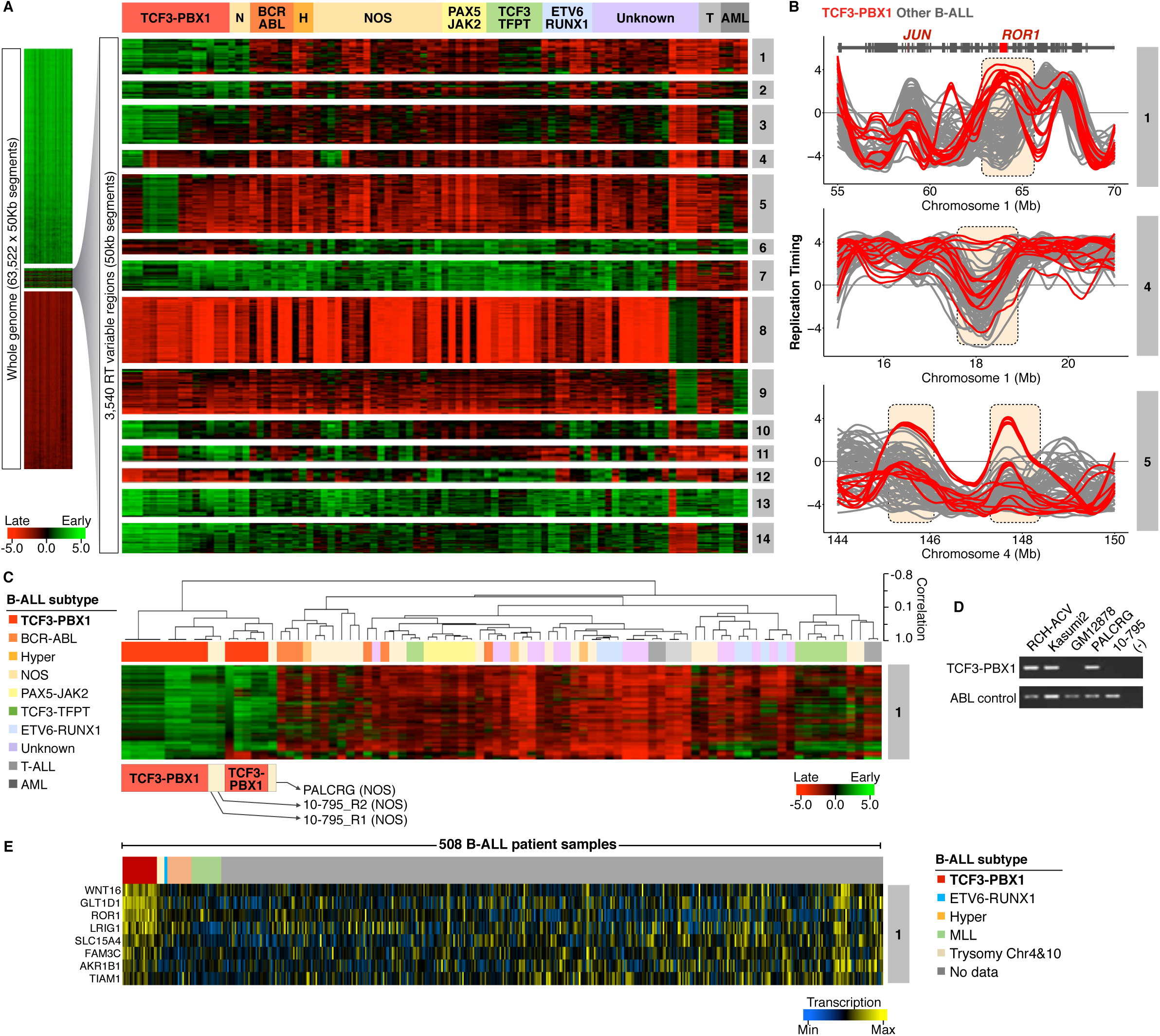
Establishing BCP-ALL RT Signatures. **(A)** patient BCP-ALL stratification using RT profiling data. RT signatures are first identified by unsupervised K-means clustering of all 50 kb genomic segments that change between early and late replication from RT log2 ratio ≥ +0.5 to/from ≤ –0.5 respectively (Rivera-Mulia et al., 2015). These stringent criteria identify features that replicate significantly different in specific clusters of patients (p-value <2×10-16 using T-tests). Branches of the dendrogram were constructed based on correlation values (distance = 1 minus the correlation value). **(B)** Exemplary RT profiles of RT signatures. The top panel show an example of an RT feature (containing the ROR1 gene) of TCF3-PBX1 (red) vs. non-TCF3-PBX1 BCP-ALL cells (grey). **(C)** Hierarchical clustering of BCP-ALL samples for RT signature #1. **(D)** RT-PCR to determine the TCF3-PBX1 fusion mRNA in patient samples classified as NOS (PALCRG and 10-795). Specific primers for the TCF3-PBX1 fusion mRNA and for ABL as a control were used. Two TCF3-PBX1-positive cell lines were used as positive controls (RCH-ACV and Kasumi2). Normal B-lymphoid cells (GM12878) was used as negative control.

Interestingly, RT signatures #1, #2 and #6 (**Figure 3A**) identified two BCP-ALLs exhibiting an RT signature of *TCF3-PBX1*-positive cells despite the samples being designated as “not otherwise specified” (NOS; cannot be classified into typical genetic subtypes including those with a t[1;19] karyotype). To determine whether these samples might have an undetected *TCF3-PBX1* fusion, we performed reverse transcriptase (RT)-PCR on mRNA isolated from these samples. Interestingly, this revealed a *TCF3-PBX1* fusion mRNA in sample PALCRG but not in sample 10-795 (**Figure 3D**), which also did not contain a *TCF3-TFPT* fusion gene (**Supplemental Table S1**) ^34^. Next, we analyzed the gene expression patterns within these RT signatures using transcriptome data from 508 B-ALL samples generated by the Therapeutically Applicable Research to Generate Effective Treatments (TARGET) from patients enrolled in Children’s Oncology Group (COG) clinical trials ^35^. This analysis identified genes with coordinated changes in transcriptional activity and RT (**Figure 3C and Supplemental Figure S3)**. Among the genes overexpressed in RT signature #1 we identified *WNT16, GLT1D1, ROR1* and others (**Figure 3D**). *WNT16* has been shown to be a target of the TCF3-PBX fusion protein and is associated with the leukemogenesis of this BCP-ALL subtype ^36^. *GLT1D* is also upregulated in *TCF3-PBX1*-positive cells but not in patients with TCF3-HLF translocation ^37^. *ROR1* encodes for the receptor orphan tyrosine kinase receptor 1 and it enhances the viability of *TCF3-PBX1*-positive cells ^38,39^. **Supplemental Table S3 contains the** complete gene list for all B-ALL RT signatures shown in **Figure 3C**.

To investigate the possible role of the *TCF3-PBX1* fusion protein in determining the RT signatures of *TCF3-PBX1*-positive leukemic cells, we transfected two *TCF3-PBX1*-positive cell lines (Kasumi and RCH-ACV) with a pair of *TCF3-PBX1*-siRNA oligonucleotides ^40^. Despite evidence of significant reduction of the TCF3-PBX1 RNA in both lines (**Figure 4A-B**), very strong genome-wide correlations in RT (≥ 0.85) were observed after downregulation of *TCF3-PBX1* (**Figure 4C-D**) and no significant changes were observed in their RT profiles (**Figure 4E-F**). Additionally, we over-expressed a TCF3-PBX1 cDNA in the mature GM12878 B-cell line (**Figure 4G**) and detected no change in its original RT profile (**Figure 4H-I**). Together, these results argue against a direct or continuing action of the *TCF3-PBX1* protein on the RT features unique to *TCF3-PBX1*-positive leukemic cells.

**Figure 4.**
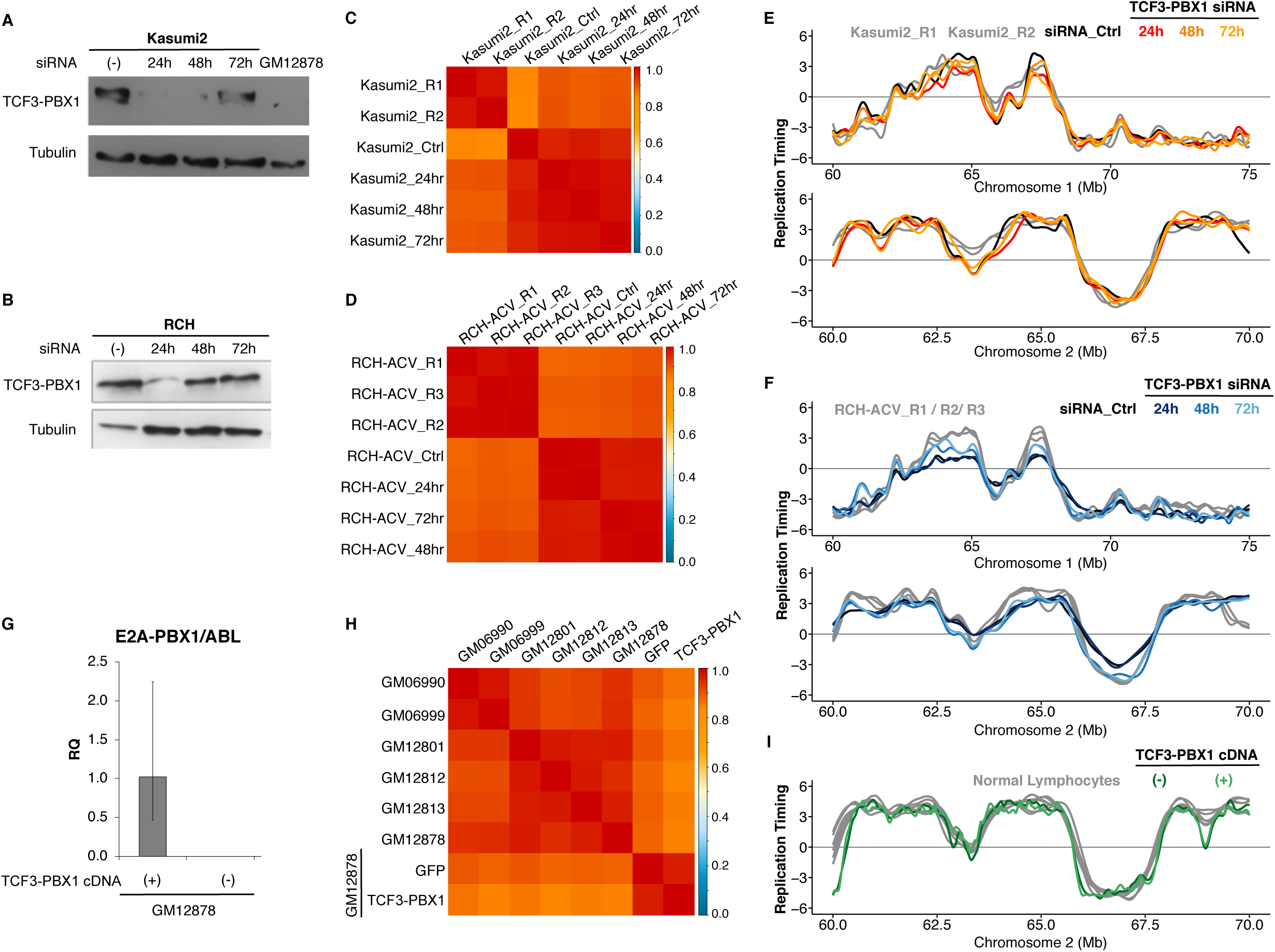
TCF3-PBX1 fusion gene does not cause the RT alterations observed in TCF3-PBX1-positive patient cells. **(A)** Down-regulation of TCF3-PBX1 fusion protein with siRNA oligonucleotides in Kasumi2 cell line. **(B)** Down-regulation of TCF3-PBX1 fusion protein with siRNA oligonucleotides in RCH cell line. **(C-D)** Genome-wide correlation matrices of Kasumi2 and RCH cells before and after down-regulation of TCF3-PBX1 fusion protein. **(E-F)** Exemplary RT profiles at two distinct chromosomal regions confirms that the RT program is preserved after down-regulation of TCF3-PBX1 fusion protein in Kasumi2 (E) and RCH (F) cell lines. **(G)** Over-expression of TCF3-PBX1 cDNA in normal B-cells (GM12878). **(H)** Genome-wide correlation matrix of normal B-cells with and without over-expression of TCF3-PBX1 cDNA. **(I)** Exemplary RT profiles confirm that the RT program is preserved after over-expression of TCF3-PBX1 fusion protein in normal B-cells.

### RT signatures associated with CNS relapse in *TCF3-PBX1*-positive BCP-ALL patients

Although most *TCF3-PBX1-positive* BCP-ALL patients are cured with current therapies, ∼10% currently relapse with a high incidence of central nervous system (CNS) relapse ^41-43^. We analyzed WGS data (28X coverage) derived from 21 *TCF3-PBX1*-positive BCP-ALL patients at diagnosis by the St. Jude’s Pediatric Cancer Genome Project (PCGP). First, we identified differences in copy number variation (CNV) and detected multiple amplified loci in all *TCF3-PBX1-positive* patients but we did not detected differences that distinguished patients with CNS relapse (**Supplemental Figure S4**). Next we exploited the deep coverage of this WGS data to generate RT profiles based on the fact that earlier replicating sequences are higher in copy number than late replicating sequences (**Figure 5A**) even without prior purification of S and G1 cells ^20^. RT datasets were derived from the 16 highest quality WGS data based on the correlation values between each other (**Supplemental Figure S4**). RT profiles across all patient samples confirmed the high concordance of the RT programs derived from WGS data (**Figure 5B**). This group of *TCF3-PBX1-positive* patients included 2 patients that had an isolated CNS relapse and 1 that developed an unrelated secondary AML. Clustering analysis of variable chromosomal segments revealed 2 RT signatures shared by the 2 CNS relapse samples (**Figure 5C**) but not the non-relapse or AML relapse samples (**Figure 5C** and **5D**).

**Figure 5.**
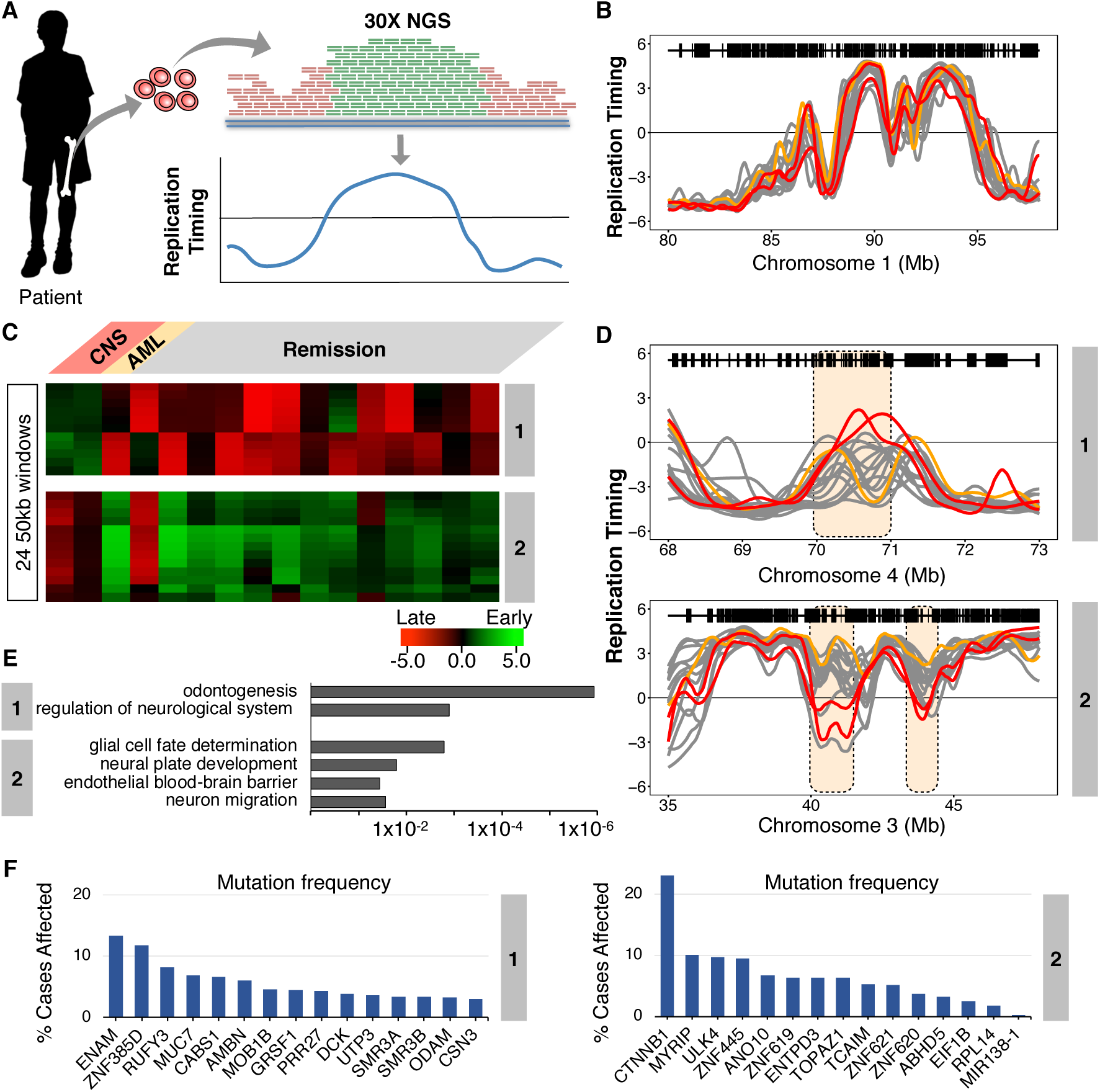
RT signatures linked to CNS relapse of TCF3-PBX1 patients. **(A)** Patient samples were subject to whole genome sequencing (WGS) at 28X depth. These sequences were subjected to a pipeline that converts their relative copy number into RT profiles (Koren et al., 2014). WGS data for 21 patients was obtained and the 16 with highest quality RT profiles (based on genome-wide correlations to each other; **Supplementary Figure S4**) were chosen for analysis. Of these, 2 samples were from patients that later suffered CNS relapse, and one was from a patient that developed a secondary AML; the remaining sampled were from patients who were in remission. **(B)** Exemplary profiles of all patient samples from a typical genomic region that shows no variance between sample profiles. **(C)** Clustering analysis of differences between relapses vs remission. 2 RT signatures distinguish patients who developed CNS relapse from those in remission. **(D)** RT profiles of each signature shown in (C). **(E)** Ontology analysis of RT signatures shown in (C). **(F)** Mutation frequencies for the genes in the RT signatures shown in (C). Mutation frequencies were obtained from the National Cancer Institute Genomic Data Commons Data Portal (Grossman et al., 2016).

RT signature #1 contained features that were early replicating in the cells from the CNS relapse samples but late in all other *TCF3-PBX1*-positive BCP-ALL patients. Gene ontology (GO) analysis of these chromosomal segments indicated that genes within this signature are associated with neurological regulation (**Figure 5E)**; specifically, this ontology term is linked to genes from the opiorphin family (*SMR3A, SMR3B* and *OPRPN*) which are up-regulated in head and neck squamous cell carcinoma (HNSCC) ^44^. Additionally, odontogenesis regulation was also associated with this RT signature, derived from genes *AMBN, AMTN, ODAM, ENAM*, which are evolutionarily related genes all located within 500 kb of each other on chromosome 4 ^45^. RT signature #2 contained features that were later replicating in samples from patients that developed CNS relapse but early in samples derived from patients in remission. GO analysis indicated that genes within this signature were associated with glial cell fate determination (Gene *CTNNB1*), neural plate development (Gene *CTNNB1*), establishment of endothelial blood-brain barrier (Gene *CTNNB1*) and regulation of neuron migration (Gene *ULK4*) (**Figure 5E**). *CTNNB1* and *ULK4* are located within 500 kb of each other on chromosome 3 (**Figure 5D)**. *CTNNB1* encodes for β-catenin, is among the most frequently mutated genes in human cancer ^46^ and has the highest mutation frequency within RT signatures #1 and #2 (**Figure 5F)**. Importantly, other genes reside in the domains with these GO-term driving genes, so further investigation is required to confirm any link between these genes and cellular phenotypes. **Supplemental Table S4** contains the complete gene list for these RT signatures.

### RT signatures associated with relapse in NOS BCP-ALL patients

A total of 120 NOS samples were obtained from the COG bank. 60 of these were from patients classified as high risk; 30 from patients that relapsed within 3 years and 30 from patients who were still in remission after at least 3 years. Because most of these samples had too few cells to support the generation of high-quality RT profiles, we selected the 5 highest quality datasets each from paired set of relapse and remission samples and generated RT signatures (**Figure 6A**). This analysis revealed 4 RT relapse-specific signatures, one of which was found in all relapses, one in 3 relapse samples and 2 in 2 relapse samples (**Figure 6B**). For all 4 signatures, all remission samples were identical. Thus, despite the high degree of heterogeneity of NOS samples, their relapse RT signatures showed several associated features. Genes found within these RT signatures are listed in **Supplemental Table S5**. GO analysis identified genes associated with interleukin-5 signaling and B-cell activation for RT signature #1; and leukocyte tethering, interleukin-35 and interleukin-21 signaling for RT signature #3 (**Figure 6C**). GO terms derived from genes *IL5RA* and *ITPR1*, which are required for B-cell development and have the highest mutation frequency within the NOS-RT signatures (**Figure 6D**).

**Figure 6.**
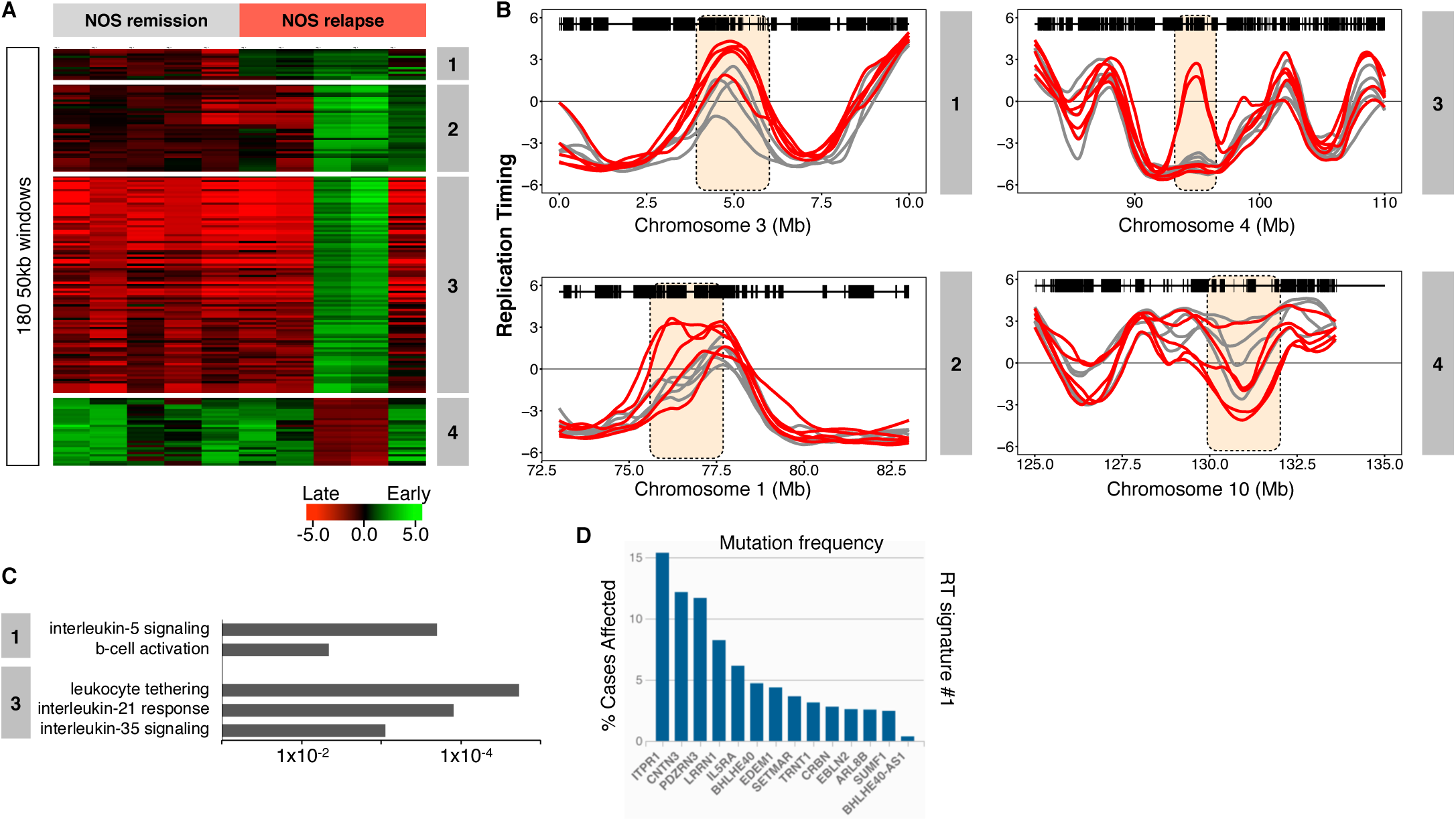
RT signatures of NOS relapse. 60 NOS samples (30 relapse; 30 remission) were obtained from the Children’s Oncology Group (COG). Many of these samples were non-viable and had to be analyzed by the repli-capture-seq S/G1 method (Figure 2), which frequently yielded a low signal to noise ratio. The 5 highest quality relapse and remission samples (based on their correlation to normal B-cells) were subjected to hierarchical clustering to identify RT signatures. **(A)** 4 RT signatures distinguish patients with relapse from those in remission. The first one includes all relapses, the 2nd only 3 relapses, and 3rd and 4th are specific for 2 relapses. **(B)** RT profiles of each signature shown in (A). **(C)** Ontology analysis of RT signatures shown in (A). **(F)** Mutation frequencies for the genes in the RT signature #1 shown in (A). Mutation frequencies were obtained from the National Cancer Institute Genomic Data Commons Data Portal (Grossman et al., 2016).

### Matching leukemia-specific RT signatures to RT profiles of normal human hematopoietic cell types

To identify features of the leukemic cell RT signatures that might match those of normal hematopoietic cells and or subsets of these at different stages of differentiation, we performed a hierarchical clustering analysis of the RT data for the subsets of genomic segments from the RT signature #1 that distinguish the *TCF3-PBX1*-positive samples including the normal blood cell types (**Figure 3A**). The genomic regions from this RT signature display heterogenous patterns of RT in mature B-cells and T-cells (**Figure 7A**). Thus, we identified sub-signatures by further dividing this subset of genomic regions by k-means clustering analysis (**Figure 7B**). This analysis identified three groups of features within signature #1, one shared with normal pre-B cells (late B-precursors) but not normal pro-B or mature B cell lines (**1A in Figure 7AB**), a second set shared with mature B-cell lines but not normal pre- or pro-B cells (**1B in Figure 7AB**), and a third set not shared with any of the nonleukemic B-cells analyzed (**1C in Figure 7AB**). When these three separate sets of features were used to subdivide signature #1, the *TCF3-PBX1* patients’ cells and cell lines, and related patient samples continued to form a closely related cohort. Additionally, *TCF3-PBX1* patient samples also cluster tightly with the RT features of the pre-B cell samples using signature #1A and with the B-cell lines using signature #1B (**Figure 7B**). Importantly, signature #1A includes the gene for the receptor orphan tyrosine kinase receptor 1 (ROR1) (**Figure 3B** and **Supplemental Table S2**). ROR1 is highly expressed in t(1;19) BCP-ALL blasts and is required for their viability, even though ROR1 is not directly regulated by either the TCF3-PBX1 fusion protein or the pre-B cell receptor (pre-BCR) ^38,39^.

**Figure 7.**
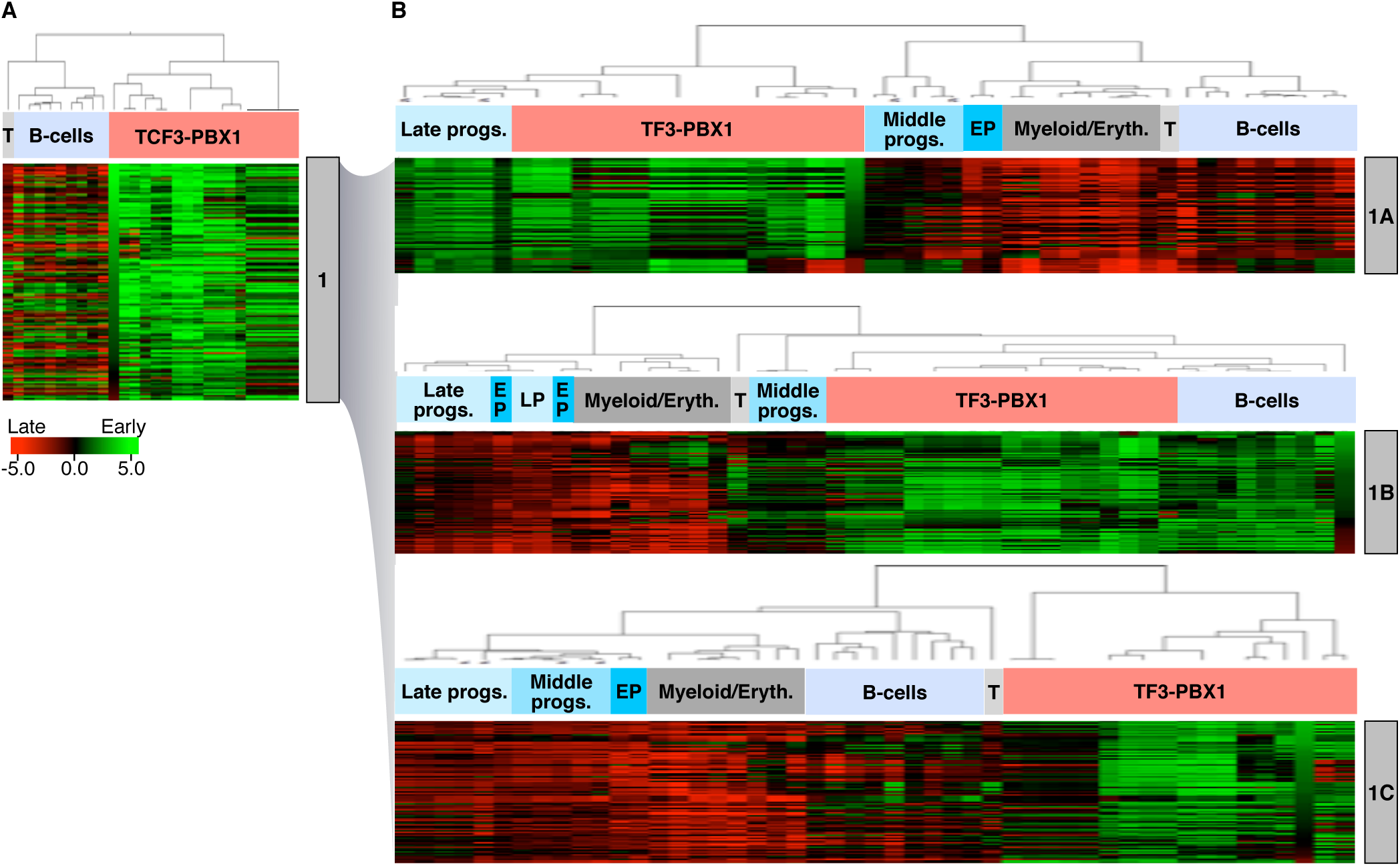
Tracking the origin of TCF3-PBX1 RT abnormalities. **(A)** Hierarchical analysis of RT signature #1 from Figure 3 clustered TCF3-PBX1 with normal B-cells. This RT signature can be sub-classified in 3 RT sub-signatures: **(B)** RT Signature #1A: TCF3-PBX1 specific RT changes that track with normal pre-B cells (late B-precursors, CD34negCD19posROR1pos cells). RT Signature #1B: TCF3-PBX1 specific RT changes that track with EBV-B cells. RT Signature #1C: TCF3-PBX1 specific RT changes that do not track with any of the B cells and substratify the patient samples.

## DISCUSSION

Our results demonstrate the potential of RT profiles to identify markers characteristic of BCP-ALL sub-types and to sub-stratify patient samples by outcome. This included the identification of several groups of domains that replicate together uniquely and consistently in TCF3-PBX1 leukemic cells as compared to other BCP-ALL subsets (**RT signatures**). Evidence that these reflect a retained epigenetic state of the normal cells they resemble is provided by the demonstration of a subset of individual domains (**RT features**) with correlates in normal CD19+CD34-pre-B cells, but not more primitive normal CD19+CD34+ pro-B cells or more mature B cells. These included ROR1, a gene that is uniquely expressed in the TCF3-PBX1 subtype of BCP-ALL where it is a therapeutic target (Bicocca, 2012). We also discovered signatures that sub-stratify TCF3-PBX1 patient samples and we identified RT signatures specific to and shared by TCF3-PBX1 as well as NOS BCP-ALL relapse cells. Other subtypes of BCP-ALL cells also contained distinguishing features. We conclude that RT profiles of BCP-ALL cells have stable prevalent clonotypic features operative in normal phenotypes they resemble as well as others with potential prognostic import.

We report several methodologies to analyze RT in normal differentiating hematopoietic cells and leukemic samples from BCP-ALL patients and to normalize them toether for comparative analyses. When possible, the method of choice is E/L Repli-seq ^17^, which yields consistently high quality data from small numbers of cells. Samples with low viability can either be be expanded as PDX to regenerate their viability ^14^ or RT profiles can be generated by a copy number comparison of reads in S-vs. G1-phase cells. Finally, when sufficiently deep WGS is available, RT profiles can be derived from total read copy number ^20^.

RT often correlates with transcription, but can clearly detect abnormalities not revealed by transcriptome analysis (Rivera-Mulia et al., 2017). The recent identification of *cis*-elements of RT control known as Early Replication Control Elements (ERCEs) reveals that RT, 3D architecture and sub-nuclear positioning are co-regulated ^47^. Thus, RT signatures, which are relatively facile to generate, have potential as a novel genre of prognostic biomarkers that reflect alterations in epigenetic regulation of large scale genome structure and function.

## ACKNOWLEDGEMENTS

We thank R. Didier for expert help with flow cytometry, and M. Hale and G. Edin and the British Columbia Cancer Agency Stem Cell Assay Laboratory staff for technical assistance in cord blood cell processing and animal irradiations. This work was funded by NIH grants R21 CA161666 and R01 GM083337, and a Margaret and Mary Pfeiffer Professorship for Cancer Research (to DG) and a Terry Fox Foundation New Frontiers Program Project grant and a Canadian Cancer Research Society Research Institute grant to CE. D.J.H.F.K. held a Canadian Institutes of Health Research (CIHR) Vanier Scholarship, and C.A.H. a CIHR Frederick Banting and Charles Best Doctoral Scholarship. Children’s Oncology Group trials were supported by grants U10 CA98543, U10 CA98413, U10 CA180886, 1U24-CA196173 and U10 CA180899 from the National Institutes of Health, and by St. Baldrick’s Foundation. The authors declare no competing financial interests.

## AUTHORSHIP CONTRIBUTIONS

J.C.R.-M., D.M.G. and C.E. designed research; J.C.R.-M., T.S., C.T.-G., N.N., D. K., C.H., K.K., V.S. and D.M.G. performed experiments; B.C., J.W.T., B.D., T.G., C.E. provided patient samples; J.C.R.-M., J.Z. and A.K. analyzed and interpreted data; J.C.R.-M. and D.M.G. wrote the manuscript with input from all authors.

## SUPPLEMENTAL MATERIAL

**Supplemental Figure S1.**
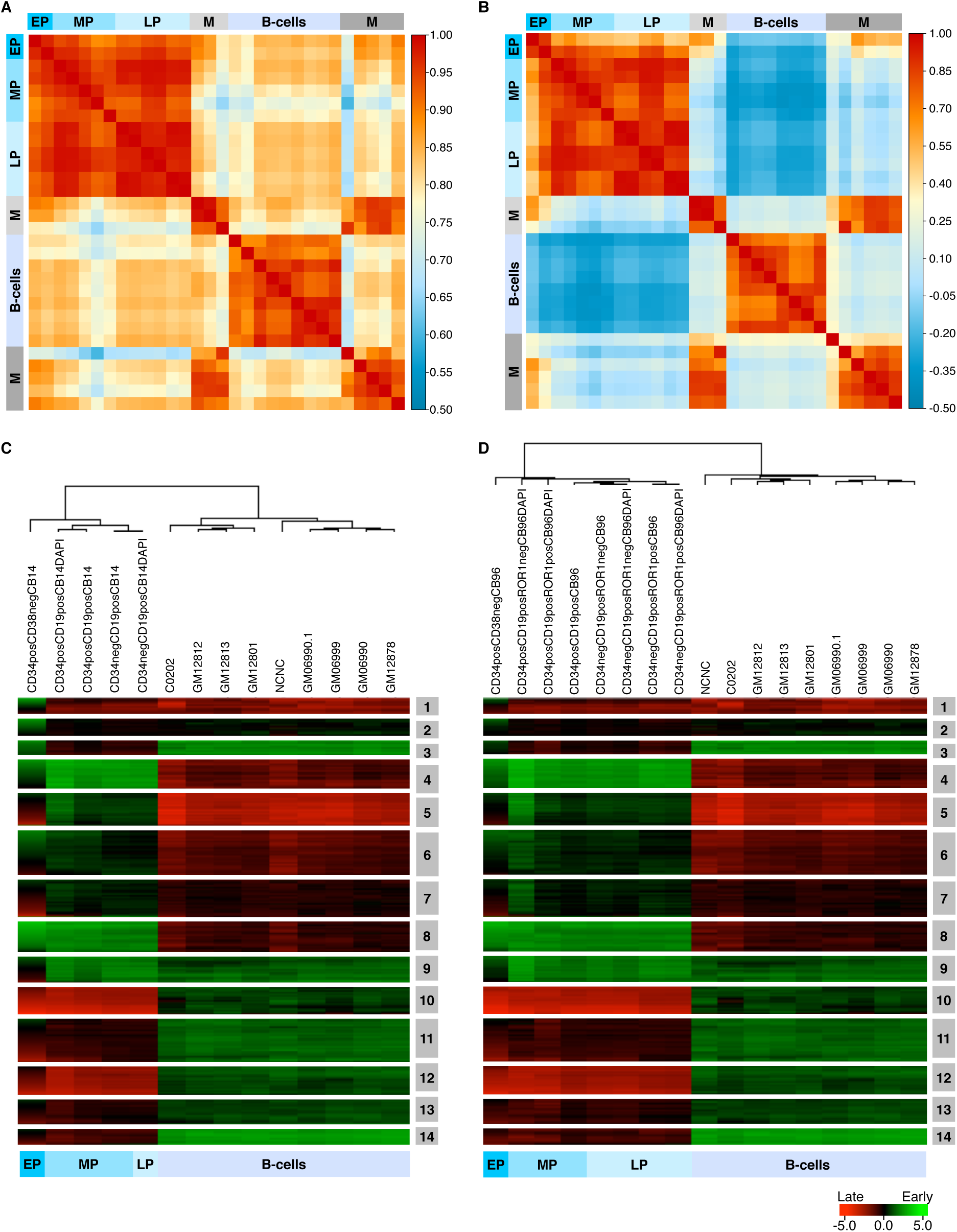
Correlation matrix of normal blood cell types. **(A)** Whole genome correlations. **(B)** Correlations using only the RT variable genome segments. **(C-D)** RT signatures per Cord Blood.

**Supplemental Figure S2.**
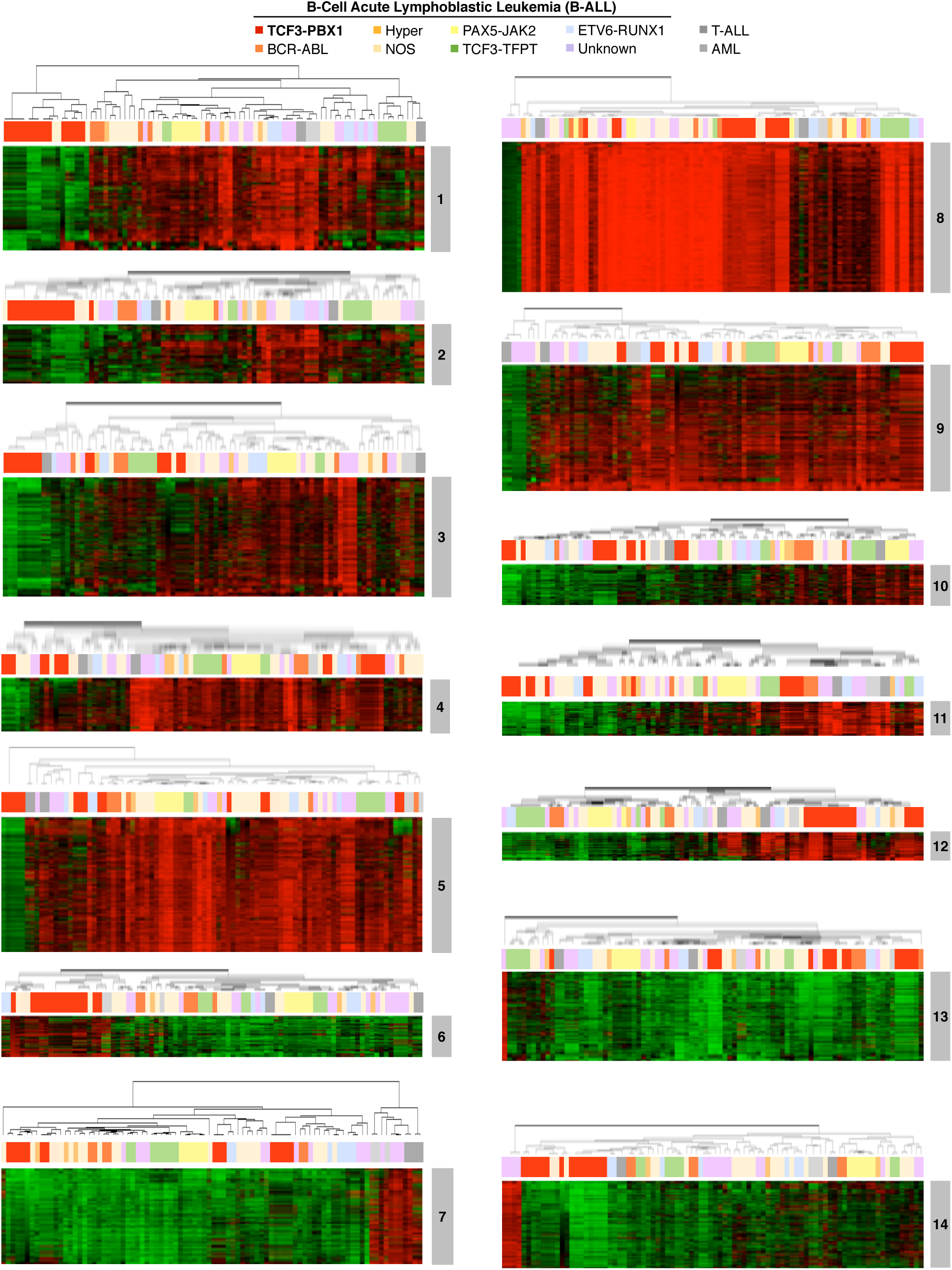
Hierarchical clustering analysis for each BCP-ALL RT signature from main Figure 3A.

**Supplemental Figure S3.**
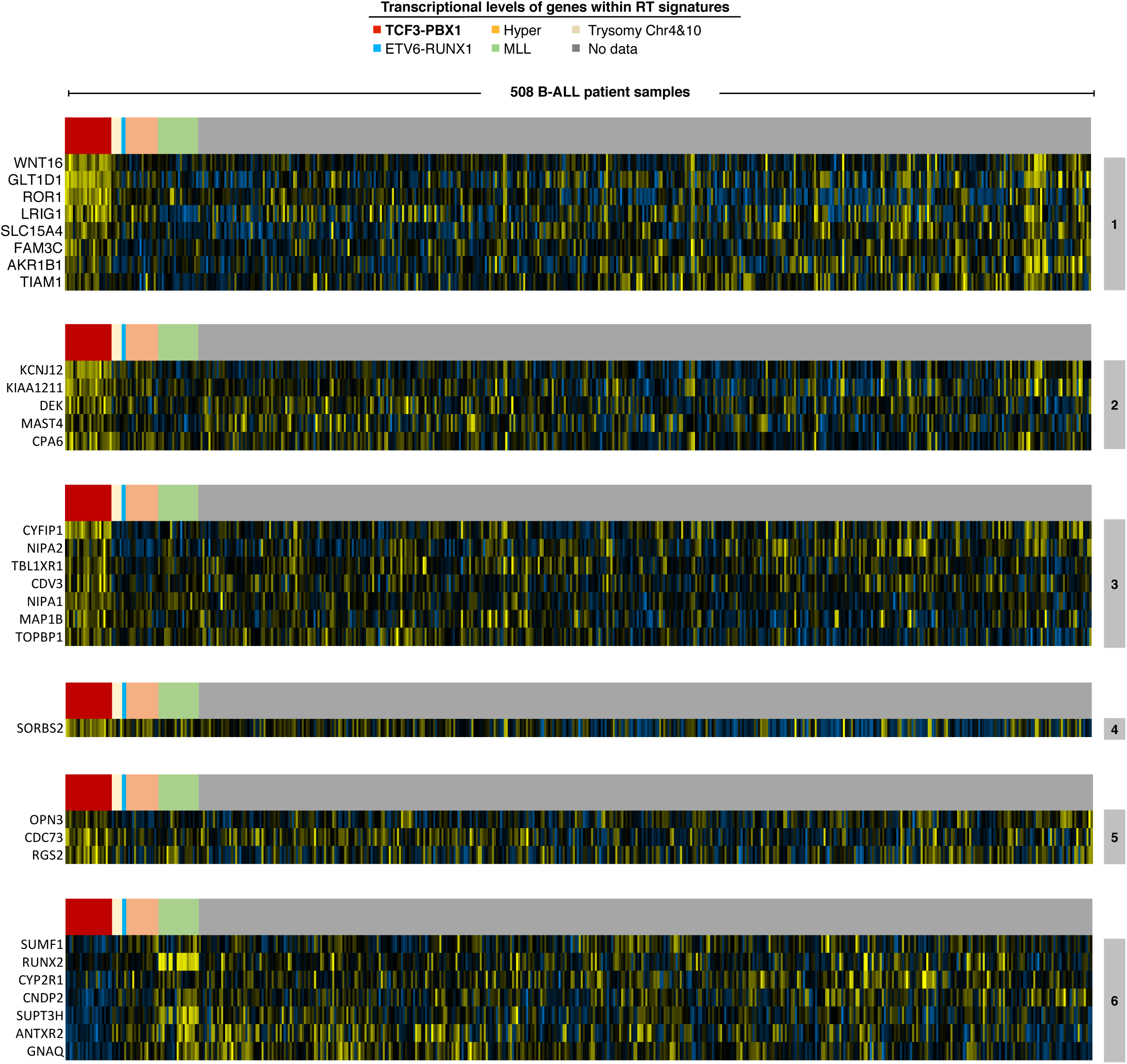
Gene expression patterns of genes within the BCP-ALL RT signature from main Figure 3A. Transcriptome data from 508 B-ALL samples was obtained from the Therapeutically Applicable Research to Generate Effective Treatments (TARGET) (Ma et al., 2018).

**Supplemental Figure S4.**
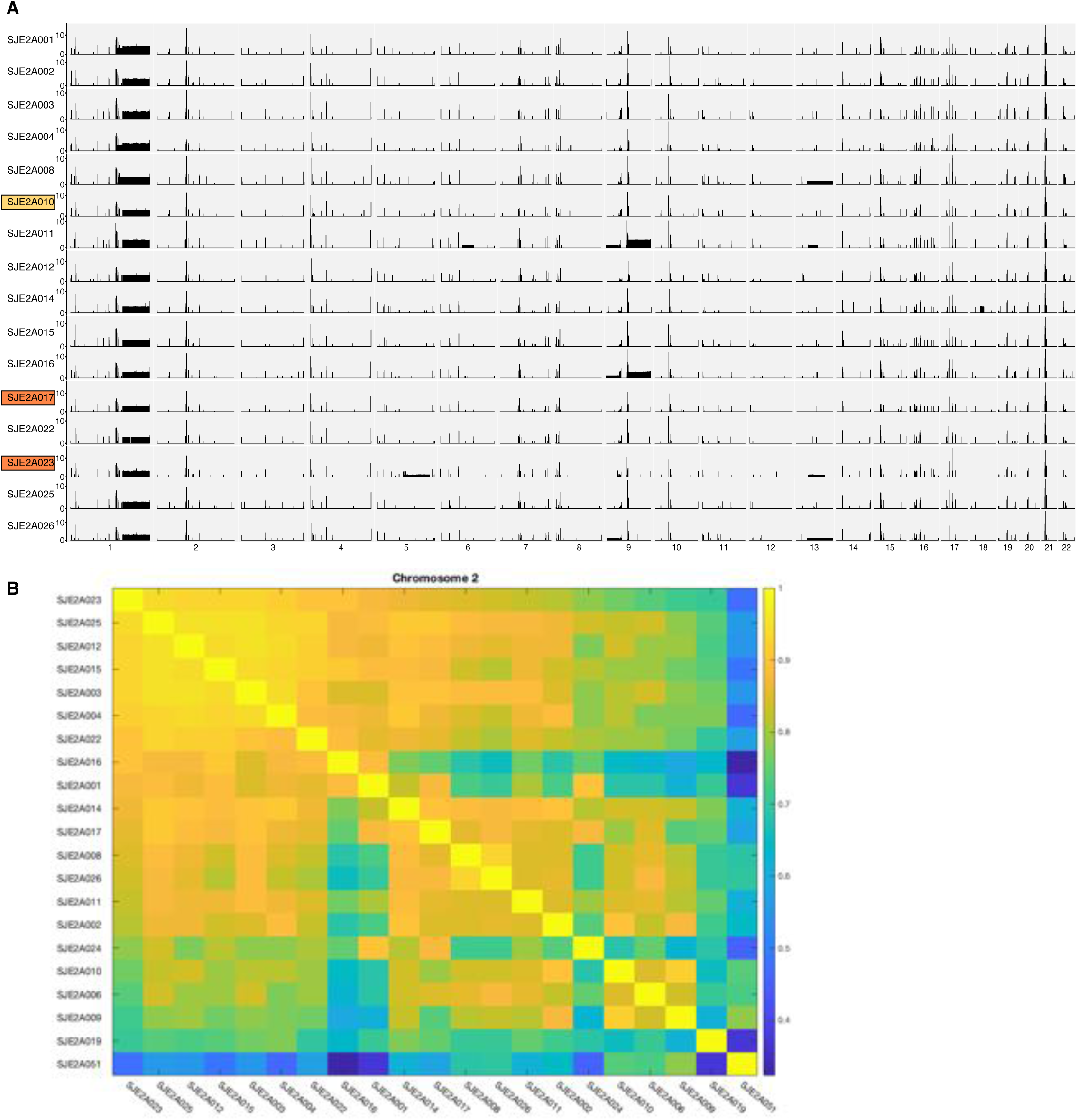
Analysis of whole genome sequencing (WGS) data (28X coverage) from TCF3-PBX1-positive BCP-ALL patients. **(A)** Copy number variation (CNV) per chromosome across TCF3-PBX1-positive patients identified multiple loci amplifications. **(B)** Correlation matrix of the RT programs derived from WGS of TCF3-PBX1-positive BCP-ALL patients. WGS data was obtained from the St. Judes Pediatric Cancer Genome Project (PCGP).

**Supplemental Table S1.** Cell samples and dataset list.

**Supplemental Table S2.** Normal blood cell types RT signatures.

**Supplemental Table S3.** B-ALL RT signatures.

**Supplemental Table S4.** TCF3_PBX1 outcome Rt signatures.

**Supplemental Table S5.** NOS outcome RT signatures.

